# Human cell-derived extracellular vesicles exhibit anti-biofilm effects against *Pseudomonas aeruginosa*

**DOI:** 10.1101/2025.07.30.667455

**Authors:** Talia J. Solomon, Casey M. Byrne, Seth S. Boledovic, Eila J. Flumen, Nicholas H. Pirolli, Emily H. Powsner, Steven M. Jay

## Abstract

The need for new antimicrobial approaches is one of the most pressing issues in modern medicine. A particular pathogen of concern is multidrug resistant *Pseudomonas aeruginosa*, which is implicated in over 1 million deaths worldwide yearly. One potential class of novel antimicrobials is extracellular vesicles (EVs), which have been found to have intrinsic antimicrobial and anti-virulence properties. Here, the antimicrobial activity of EVs on *P. aeruginosa* was explored in the context of biofilms of hyper-virulent strain PA14. We identified the human monocyte cell line THP-1 as a promising source of EVs for this application, inducing reduced PA14 biofilm formation in a dose-dependent manner. THP-1 EVs were not found to affect *P. aeruginosa* growth planktonically within the biofilm assay or in shaken culture. Additionally, we demonstrate that anti-biofilm effects were conserved with similar efficacy across THP-1 monocyte and differentiated THP-1 macrophage-derived EVs. Further, EVs from induced pluripotent stem cell-derived mesenchymal stem/stromal cells (iMSCs) also reduced PA14 biofilm formation to a similar extent as THP-1 EVs, while EVs from human embryonic kidney cells (HEK293T) had a similar effect to its media control. This work indicates that human cell-derived EVs from several sources possess biological and/or physical properties that reduce PA14 biofilm formation.

## INTRODUCTION

Antimicrobial resistance is implicated in over 35,000 deaths in the US annually (2019) and 1.27 million deaths worldwide (Murray 2022), with these numbers only expected to increase in the future. In particular, *Pseudomonas aeruginosa* has been identified as a serious threat by the CDC and a critical first priority pathogen by the WHO (2017, 2019). As clinically significant resistance has been observed against every antibiotic in use (Clatworthy, Pierson et al. 2007, Dickey, Cheung et al. 2017), the need for new antimicrobial approaches against *P. aeruginosa* and other pathogens is urgent and clear. As such, there has been increased pursuit of novel antimicrobial strategies, including phage therapy, immunomodulation, antisense RNA, antimicrobial peptides (AMPs), and antibodies (Baker, Payne et al. 2018, Theuretzbacher and Piddock 2019, Theuretzbacher, Outterson et al. 2020). Some of these are direct-acting therapeutics, meaning that they inhibit growth or kill microbes, whereas others aim to alter the immune response to the pathogen or to specifically block a virulence factor of the pathogen (Theuretzbacher, Outterson et al. 2020). Anti-virulence treatments are exciting because they have the potential to impose less evolutionary pressure toward the development of resistance and may have a decreased effect on the microbiome (Clatworthy, Pierson et al. 2007, Dickey, Cheung et al. 2017, Fleitas Martínez, Cardoso et al. 2019, Monserrat-Martinez, Gambin et al. 2019). However, it remains unclear whether they actually will contribute to less resistance, and there is debate over whether combination therapy with direct-acting medications would be necessary to kill the microbial infection (Maura, Ballok et al. 2016, Theuretzbacher, Outterson et al. 2020). Additionally, phage therapy and some of the anti-virulence therapeutics are very strain-specific, significantly limiting the spectrum of activity and utility (Theuretzbacher and Piddock 2019). Furthermore, there are challenges with the pharmacokinetics of RNA- and protein-based therapeutics (Mahlapuu, Håkansson et al. 2016).

Alternatively, extracellular vesicles (EVs) represent an emerging modality that can potentially augment the antimicrobial arsenal. EVs have been shown to have inherent antimicrobial and anti-virulence properties (Schulz, Goes et al. 2018, Shopova, Belyaev et al. 2020, Koeppen, Nymon et al. 2021, Zhao, Zhang et al. 2022) in addition to being natural nanoscale delivery vehicles that are capable of carrying RNA- and protein-based therapeutics (Wiklander, Brennan et al. 2019). Antimicrobial EVs have been derived from a variety of cell types, including epithelial cells of the lungs (Koeppen, Nymon et al. 2021), kidney (Hiemstra, Charles et al. 2014), and oral mucosa (Koeppen, Nymon et al. 2021, Zhao, Zhang et al. 2022) as well as mesenchymal stromal/stem cells (MSCs) (Monsel, Zhu et al. 2015, Alcayaga-Miranda, Cuenca et al. 2017, Jonnalagadda 2021), and immune cells like neutrophils (Shopova, Belyaev et al. 2020). EVs have been shown to have activity against a variety of pathogenic microbes, including *P. aeruginosa* (Brakhage, Zimmermann et al. 2021, Koeppen, Nymon et al. 2021), and other bacteria such as *Escherichia coli* (Hiemstra, Charles et al. 2014, Monsel, Zhu et al. 2015, Schulz, Goes et al. 2018, Brakhage, Zimmermann et al. 2021, Jonnalagadda 2021), *Staphylococcus aureus* (Keller, Ching et al. 2020, Brakhage, Zimmermann et al. 2021, Jonnalagadda 2021), and *Klebsiella pneumoniae* (Jonnalagadda 2021), as well as fungal pathogens like *Aspergillus fumigatus* (Shopova, Belyaev et al. 2020) and *Candida albicans* (Zhao, Zhang et al. 2022). With regard to *P. aeruginosa*, the work of Koeppen et al. showed that airway epithelial cell (AEC)-derived EVs inhibit strain PA14 biofilm formation via miRNA let-7b-5p transfer (Koeppen, Nymon et al. 2021). Additionally Hendricks et al. demonstrated that AEC-derived EVs from cells infected with RSV led to enhanced *P. aeruginosa* biofilm formation due to the transfer of nutrient iron via transferrin (Hendricks, Lane et al. 2021). These studies lay the framework for a more detailed analysis of the potential of human cell-derived EVs as antimicrobial effectors against *P. aeruginosa*.

Following from the works cited above, here we studied the potential of EVs from the monocyte-like cell line THP-1 to reduce biofilm formation of *P. aeruginosa* strain PA14, a commonly-used reference strain that models a hyper-virulent phenotype (Rahme, Stevens et al. 1995). THP-1 cells were chosen as EV producers based on prior demonstrations of the potential of macrophage and monocyte EVs as inter-domain effectors (Ding, Rivera et al. 2020, Halder, Babych et al. 2022) as well as the increased potential of a cell line such as THP-1 to be engineered and used for scalable EV manufacturing compared to primary cells such as bone marrow-derived monocytes. The ability to disrupt biofilms was focused on due to its relevance in potential therapeutic applications for diseases such as cystic fibrosis and others (Reynolds and Kollef 2021).

## MATERIALS AND METHODS

### Cell culture

THP-1 monocytes (ATCC TIB-202) were cultured in RPMI media with 10% FBS and 1% Penicillin/Streptomycin and passaged every 3-5 days. Cells were maintained at concentrations between 100,000 and 1,000,000 cells/mL. To prepare cells for EV collection, cells were spun at 220xg for 5 minutes and then the media was changed to EV depleted RPMI media. This media was prepared as RPMI media with 10% FBS (no antibiotics were used in EV depleted media except for in experiments displayed in Figure 4) and then the media was run through the tangential flow filtration system to remove particles larger than 100 kDa. Media was EV depleted as whole media as opposed to EV depletion of FBS alone to minimize effects of media on functional experiments (Kronstadt, Van Heyningen et al. 2023). iMSCs were cultured in DMEM supplemented with 10% FBS and 1% Penicillin/Streptomycin and MEM Non-Essential Amino Acids. HEK293T cells were cultured in DMEM supplemented with 10% FBS and 1% Penicillin/Streptomycin.

For THP-1 differentiation, 40 million THP-1 cells were placed into a new flask and 100 uM PMA was spiked into the flask to differentiate the THP-1 monocytes into macrophages. Following differentiation into macrophages, the media was replaced with either M1 or M2 differentiation media. For M1 differentiation, media was RPMI with 20 ng/mL IFNg and 10 pg/mL LPS. For M2 differentiation, RPMI was supplemented with 20 ng/mL IL4 and IL13 (Genin, Clement et al. 2015, Scheurlen, Snook et al. 2021). After cells were differentiated to the desired state, the cells were washed with PBS and media was replaced with EV depleted RPMI and conditioned media was collected each day for the following 2-3 days post differentiation.

*Pseudomonas aeruginosa* strains PA14(Rahme, Stevens et al. 1995) were maintained in a 10% glycerol stock stored at −80 °C. Prior to an experiment, the glycerol stocks were streaked onto LB Miller agar (1%) plates and incubated at 37 °C overnight. The following day, a colony was selected and used to inoculate a 5 mL culture in LB miller broth which was then incubated shaking at 37 °C.

### EV isolation

EV isolation was completed as previously described (Levy, Abadchi et al. 2023). THP-1 EVs were generally isolated from volumes ranging from 150-300 mL of conditioned media and 70-100 mL of conditioned media for differentiated THP-1 EVs. To isolate THP-1 EVs, cells were maintained at 600,000 cells/mL each day of collection. Cells were diluted to this concentration daily and then at the end of collection, the cells were pelleted at 220xg for 5 minutes. Following removal of cells, the media underwent three differential spins. First the conditioned media was spun at 1,000xg for 10 minutes and then the media was transferred to new tubes leaving a few mL of media with pelleted cells and debris behind at the bottom of each tube. Similarly, the conditioned media was then spun at 2,000xg for 20 minutes and then transferred to new tubes and then spun at 10,000xg for 30 minutes. Following this spin the conditioned media was filtered through a 0.22 um PES sterile filter and then tangential flow filtration was used to concentrate and buffer exchange to PBS. A 100 kDa molecular weight cutoff (MWCO), modified PES, membrane was used in the tangential flow filtration process (Busatto, Vilanilam et al. 2018). After the sample was collected from the tangential flow filtration process, it was concentrated in 100 kDa MWCO PES centrifugal filters to approximately 0.2 mL. The following sample was resuspended at approximately 400 uL total volume and sterile filtered through a 0.22 um PES sterile syringe filter. EVs were stored at −20°C and used within 4 weeks of isolation (Levy, Jeyaram et al. 2023).

### EV characterization

Nanosight (Malvern Technologies) was utilized to quantify the EV concentration and diameter in each EV sample. The EV sample was diluted appropriately (approximately 1:100 dilution) in PBS and then analyzed on the Nanosight utilizing a camera level of 7 and threshold of 3. EV protein content was analyzed with a BCA assay per manufacturers instructions (G Bioscience). Isolated EVs were used directly or diluted 1:10 to assess protein content. 25 uL of EV dilutions were plated along with 25 uL of the BSA protein standards. 200 uL of BCA working reagent and incubated at 37°C for 30 minutes. Absorbance was measured on a plate reader (Spark, Tecan).

### Bacterial growth assay

To study the effects of EV treatments on growth, a growth curve assay was performed utilizing a Tecan Spark plate reader. Prior to the experiment, *P. aeruginosa* strain PA14 was streaked on an agar plate and a single colony was used to start an overnight culture at 37°C. Then the optical density of the overnight culture was measured and it was diluted to OD600 of 0.1. Then the subculture was incubated at 37°C until the culture was in log phase (OD600 0.5-2) which took approximately 3 hours. Meanwhile, a 96 well plate was prepared for the assay, diluted EV treatments and controls were plated each 100 uL into 3 wells. EV treatments were at a concentration of 5*10^9 EVs/mL unless otherwise stated (Koeppen, Nymon et al. 2021). Additionally, 200 uL filtered water was plated into the wells along all of the edges of the plates to provide humidity to limit evaporation losses (Schulz, Goes et al. 2018). Once the treatments and water were plated, 100 uL of the *P. aeruginosa* subculture dilution was plated, this was diluted to OD600 0.02. Then the plate was incubated in the Tecan, Spark multimode plate reader with humidity cassette. 4 mL of filtered water was placed in each side of the humidity cassette and the plate was incubated inside of the humidity cassette without the 96 well plate lid, lid utilizing the magnetic lid of the humidity cassette. OD600 readings were taken every 15 minutes overnight for approximately 16 hours. Note, for this assay, treatment groups were placed in columns and shuffled around the plate for each replicate to control for well effects.

### Biofilm assay

To assess EV activity on *P. aeruginosa* biofilms, the general protocol of O’Toole was followed (Au - O’Toole 2011). Briefly, prior to the experiment, *P. aeruginosa* was streaked and cultured overnight. Then the overnight cultures were diluted 1:100 with fresh LB broth. Meanwhile, a 96 well plate was prepared for the assay, diluted EV treatments and controls were plated each 100 uL into 3 wells. EV treatments were at a concentration of 5*10^9 EVs/mL unless otherwise stated (Koeppen, Nymon et al. 2021). Additionally, 200 uL filtered water was plated into the wells along all of the edges of the plates to provide humidity to limit evaporation losses. Once the treatments and water were plated, 100 uL of the *P. aeruginosa* dilution was plated. The plates were then left to incubate at 37 °C for 24 hours. A beaker of water was left in the incubator to provide more humidity to prevent evaporation losses. Following the incubation, the OD600 was measured using a Spark Tecan plate reader. Then the planktonic culture was dumped out and the plate and adhered biofilms were washed 2x in filtered water and then blotted dry onto paper towels. 250 uL of 0.1% crystal violet was pipetted into each well and allowed to incubate for 15 minutes at room temperature. Following this the excess dye was dumped out and then the plate was washed and blotted 4x. The plate was then allowed to dry overnight before quantification. 30% acetic acid was used to dissolve the crystal violet that stained the biofilm. To do so, 250 uL was pipetted into each well and then the plate was incubated rocking for 15 minutes at room temperature. Then 200 uL of the mixture was pipetted into a new plate and the absorbance at 550 nm was measured using a Spark Tecan plate reader. 30% acetic acid was used as a blank measurement for the data analysis. Note, for this assay, treatment groups were placed in columns and shuffled around the plate for each replicate to control for well effects.

### Statistical analysis

Data are presented as mean +/− standard error of the mean. An ordinary one-way ANOVA was performed with either a Sidak’s or Tukey’s multiple comparison test. Statistical analyses were performed with Prism 10 (GraphPad Software). Statistical significance is shown as *p<0.05, **p<0.01, ***p<0.001, ****p<0.0001.

## RESULTS AND DISCUSSION

### THP-1 EVs inhibit PA14 biofilm formation

To explore the antimicrobial potential of THP-1 EVs against *P. aeruginosa*, THP-1 EVs were isolated and characterized (Figure S1) before being utilized in PA14 growth and biofilm formation assays. A dose of 5E9 EVs/mL (as determined by NTA) was selected as an initial dosage to study based on literature (Koeppen, Nymon et al. 2021) and prior experience from other functional assays (Levy, Abadchi et al. 2023). This dose of THP-1 EVs was found to reduce PA14 biofilm formation over a 24-hour period compared to control (Figure 1A, E). Additionally, this effect was found to be dose-dependent in a crystal violet microtiter dish biofilm formation assay (Figure 1B). Interestingly, THP-1 EV treatments had no observable preventative effect on PA14 growth in suspension (Figure 1C-D,F), suggesting that the anti-biofilm effect is not related to decreasing PA14 growth and classifying THP-1 EVs as potential anti-virulence, anti-biofilm therapeutics. These results are similar to those found by Koeppen et al. for AEC EVs (Koeppen, Nymon et al. 2021), demonstrating a partial prevention in biofilm formation without inhibiting planktonic growth of PA14 cells. Contrastingly, work from the Morici lab has demonstrated that bacterial EVs (BEVs) from *Burkholderia thailandensis* have an antimicrobial effect against several organisms including *P. aeruginosa*, with an anti-biofilm effect against *Streptococcus mutans* due to specific antimicrobial compounds that limit pathogen growth and thus biofilm formation (Wang, Hoffmann et al. 2020, Wang, Hoffmann et al. 2020).

**Figure 1.**
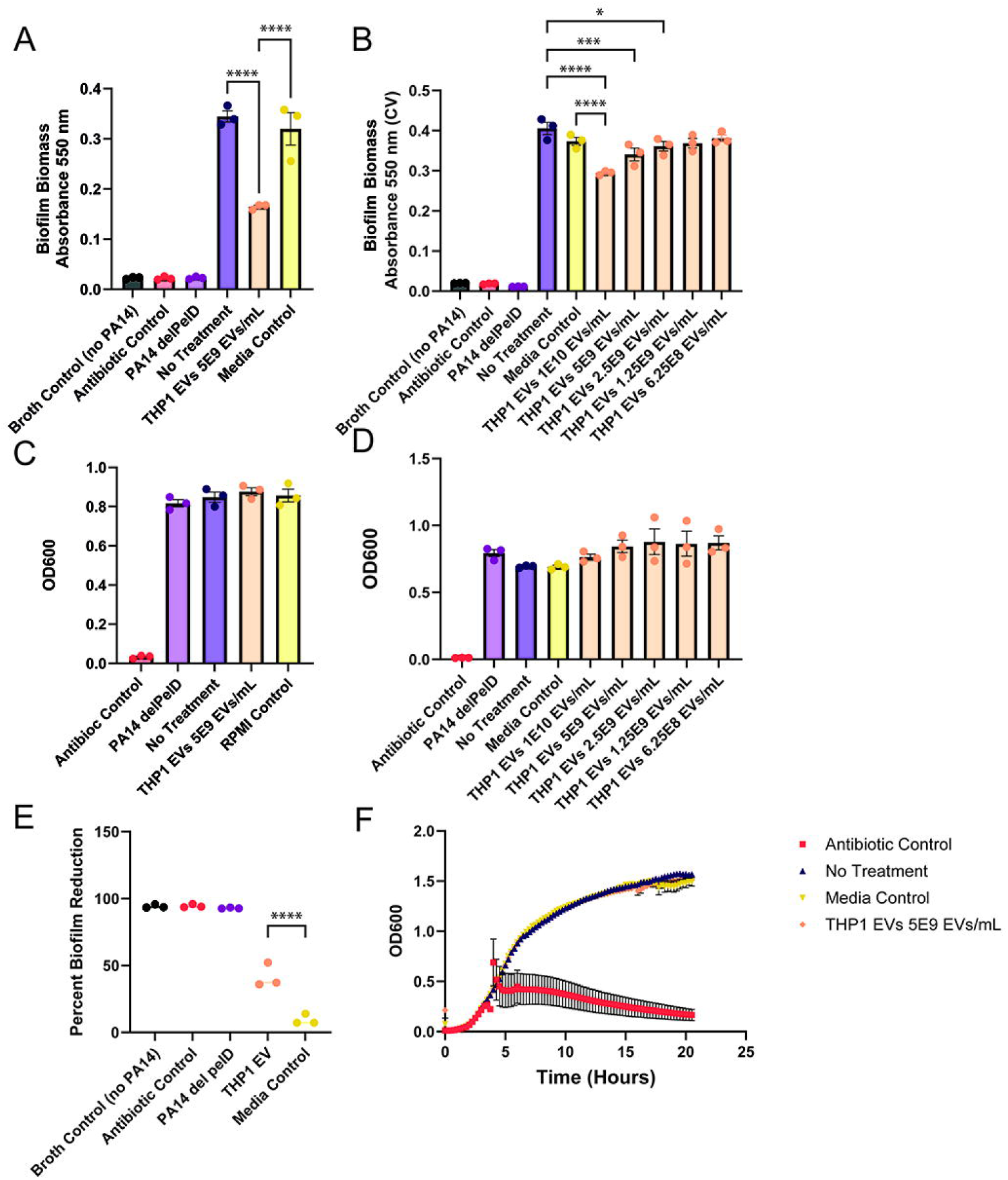

Of note, prior studies from our research group have revealed confounding effects of media contaminants in EV batches on the results of functional assays (Kronstadt, Van Heyningen et al. 2023) and so an additional media control was employed to help distinguish between the effects of media contaminants from the isolated vesicles. This media control was EV depleted RPMI media (supplemented with FBS) that underwent the EV isolation procedure and was dosed in equal volume as identically-processed EV samples. We found that the media controls depending on the batch have a small ability to prevent biofilm formation, and so these controls were used in all experiments presented here. To confirm whether intact THP-1 EVs were necessary for this anti-biofilm effect, THP-1 EVs were incubated at room temperature (RT) for 2 weeks to allow for degradation of the vesicles (Lee, Ban et al. 2016) and then tested in the biofilm formation assay. RT-incubated vesicles were ineffective at preventing PA14 biofilms as compared to no treatment control (Figure 2A, C), and both the RT-vesicles and RT-incubated media control induced an increase in biofilm biomass without a corresponding increase in planktonic PA14 growth (Figure 2B), again indicative of a differential interaction with THP-1 EVs between planktonic and biofilm forms of PA14. This last result suggests that loss of anti-biofilm activity due to degraded EV structure could be further amplified by potential utilization of degraded EVs as nutrients, echoing the work by Hendriks et al.(Hendricks, Lane et al. 2021), or as biofilm matrix components, as seen with *P. aeruginosa*-derived EVs (Schooling and Beveridge 2006).

**Figure 2.**
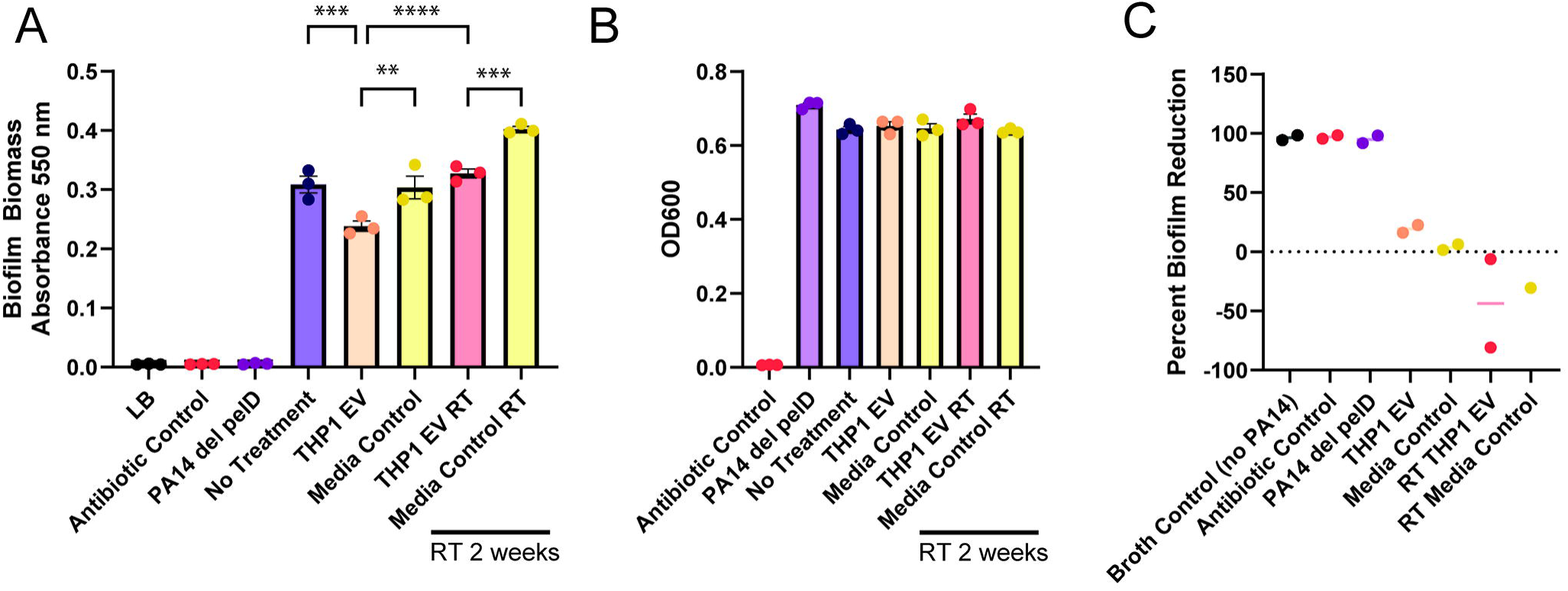

### Differentiation of THP-1 source cells does not change anti-biofilm effect of EVs

To assess potential mechanisms of the anti-biofilm effect of the THP-1 EVs, THP-1 monocytes were differentiated into a basal macrophage (M0), inflammatory macrophage (M1), and a regenerative macrophage (M2) states prior to EV collection to assess any changes in anti-biofilm effect of their EVs. After differentiation to basal macrophages, cellular morphology shifted to an adherent cell type that was confirmed visually. Further, macrophage polarization was assessed with qPCR for macrophage markers, which showed enhanced TNFa expression in naïve macrophages compared to THP-1 monocytes, with even higher expression in inflammatory macrophages (Figure S2A). Additionally, regenerative macrophages had increased CD206 expression (Figure S2B). Interestingly, the naïve and inflammatory macrophages had higher IL10 expression than the regenerative macrophages after polarization, though it was not statistically significant due to variability across biological replicates (Figure S2C). When the EVs of these differentiated THP-1 cells were tested in the biofilm formation assay, there were no statistically significant differences observed between experimental groups, while the general anti-biofilm effect was retained (Figure 3A-B). As in the prior experiments, no effect of any EV group on planktonic growth of PA14 cells was seen (Figure 3C). These results were surprising, as the contents of EVs are determined by the phenotype of their source cells (Wiklander, Brennan et al. 2019) and other work has shown that EV content and function changes upon polarization of source macrophages (Bouchareychas, Duong et al. 2020). We had expected that the inflammatory or “M1” macrophage-derived EVs would have the most potent anti-biofilm effect since they are most associated with infection fighting and would thus potentially have more anti-microbial cargo, but this was not the case.

**Figure 3.**
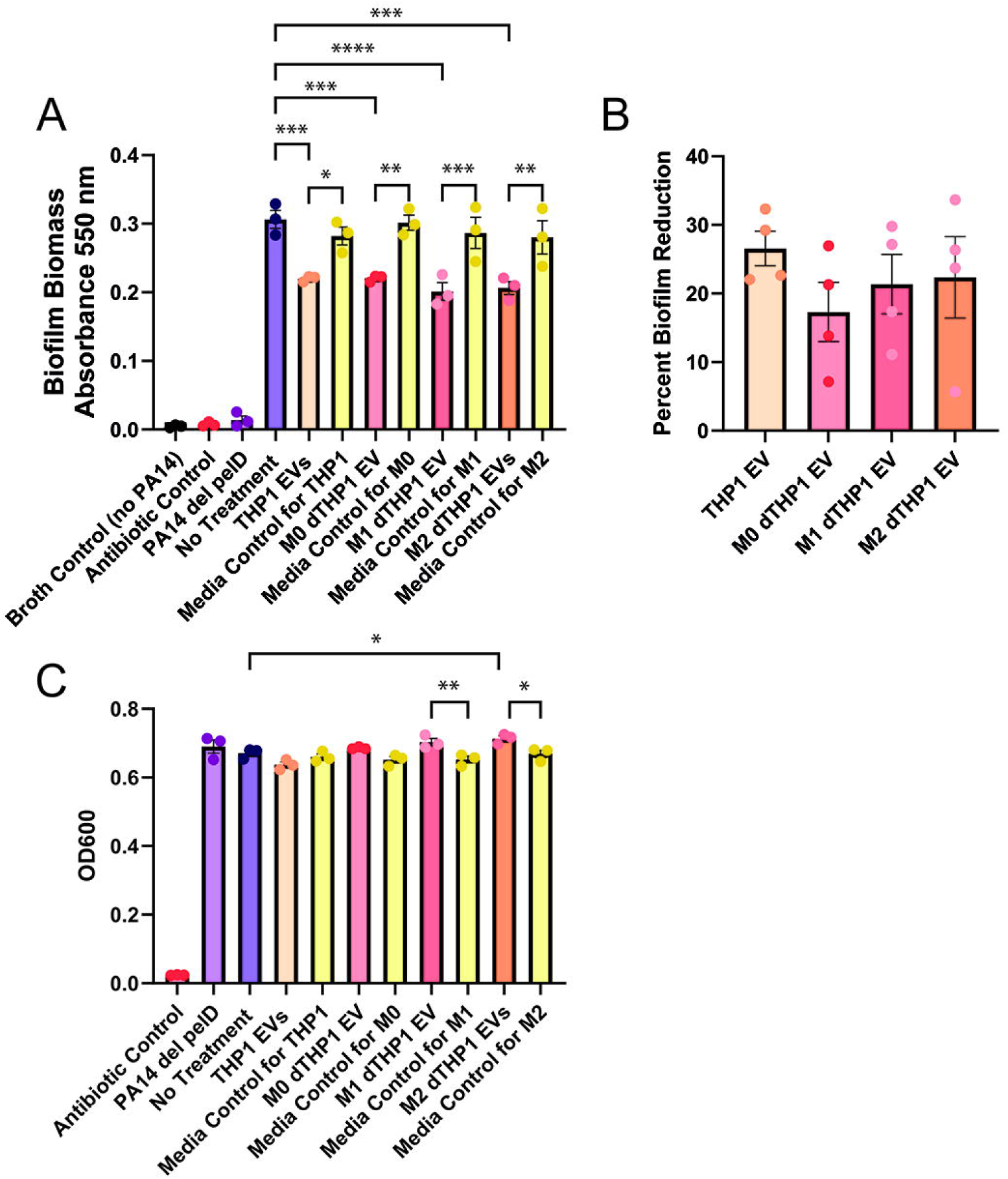

### Specificity of human cell-derived EVs on PA14 biofilm inhibition

Finally, since all viable EV groups tested so far had exhibited anti-biofilm effects, we tested additional EV sources to determine if this might be a non-specific effect of human cell-derived EVs. We chose to test commonly-used human EV sources: induced pluripotent stem cell-derived mesenchymal stem/stromal cells (iMSCs) and human embryonic kidney cells (HEK293Ts). HEK293T cells are popularly used for genetic engineering as they transfect easily, while iMSCs are favored in EV research for regenerative and anti-inflammatory purposes and have been studied in infection models (Alcayaga-Miranda, Cuenca et al. 2017, Yoon, Lee et al. 2023). In line with prior data, iMSC EVs demonstrated an anti-biofilm ability compared to the no treatment and media controls (Figure 4A, E) while not decreasing growth of PA14 planktonically (Figure 4C), similar to THP-1 EVs. On the other hand, HEK293T EVs showed a limited ability to effect biofilm formation that was similar to that of the media control (Figure 4B, F), where in one replicate both the media control and HEK293T EVs decreased biofilm formation a little and in the other they both slightly increased biofilm formation. This is in line with our group’s prior findings regarding the effect of media contaminants confounding the effect of HEK293T EVs (Kronstadt, Van Heyningen et al. 2023). Neither of these EV populations decreased PA14 growth planktonically (Figure 4D). These data suggest some specificity of the anti-biofilm effects of THP-1 and iMSC EVs, possibly due to their molecular cargo, but also leave open the possibility that some conserved factor or factors in EVs generally reduce PA14 biofilm formation in this assay. The latter could be due to the presence of lipids or perhaps the physical displacement of bacteria via EVs during biofilm initiation.

In conclusion, we report that THP-1 EVs do not impact the growth of *P. aeruginosa* strain PA14, but do reduce biofilm formation in a microtiter dish assay, demonstrating partial prevention of biofilm formation on a synthetic surface. We further observed that the anti-biofilm effect was not universal among human cell-derived EVs, suggesting a degree of specificity in the interaction between THP-1 EVs and *P. aeruginosa*. However, the anti-biofilm effect was retained when THP-1 cells were differentiated into macrophages in naïve (M0) and polarized (M1, M2) phenotypes, and other human cell-derived (iMSC) EVs also showed anti-biofilm effects. These results suggest potential roles for both EV molecular cargo as well as structural or physical aspects of EVs, in the reduced biofilm formation observed, such as EVs disrupting the interaction between PA14 and its growth substrate. Overall, this work provides further insight into the role human cell-derived EVs can play in biofilm prevention and furthers knowledge towards exploration of EVs as anti-biofilm therapeutics.

**Figure 4.**
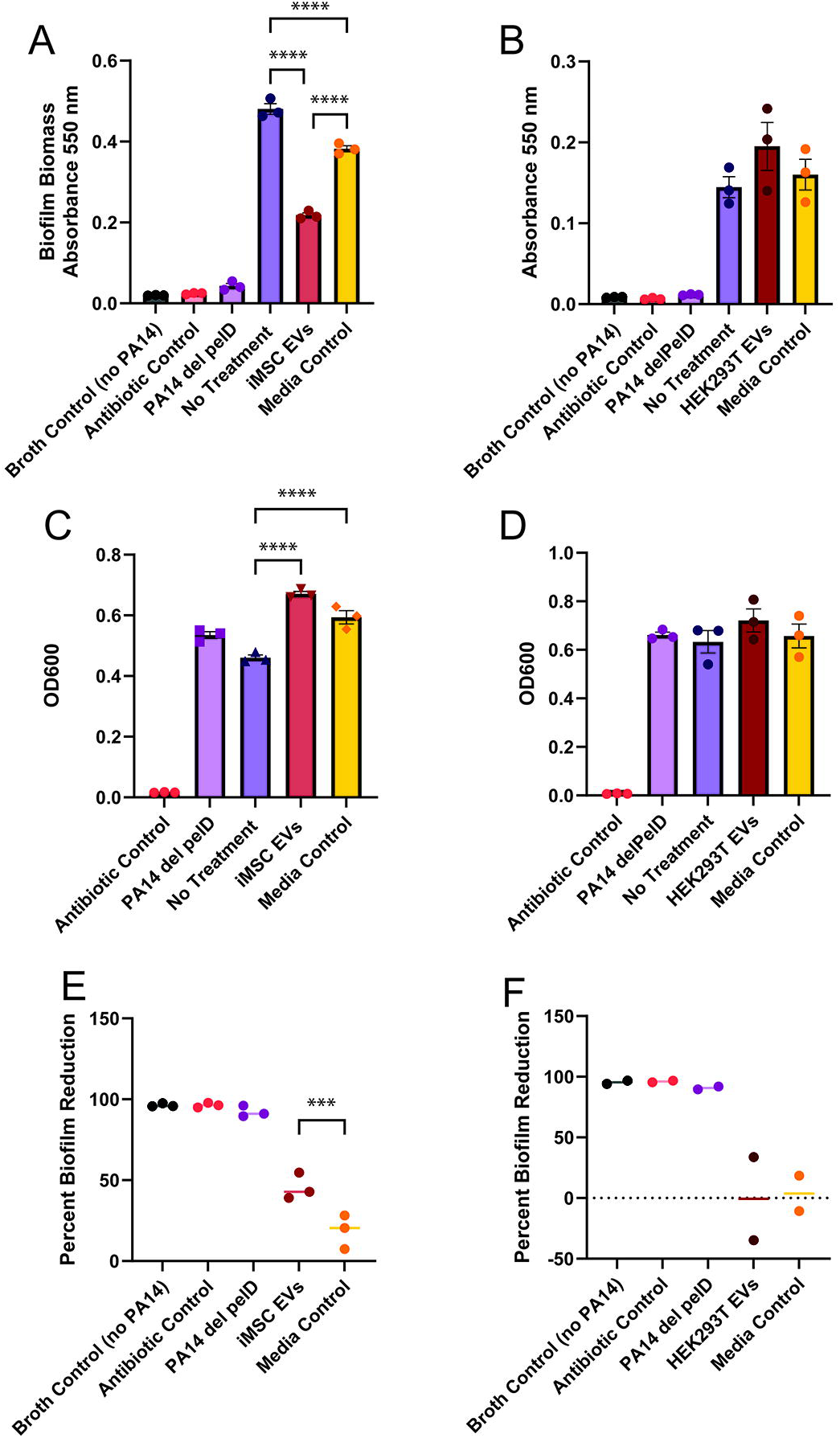

