## Supplementary Material for "Human cell-derived extracellular vesicles exhibit anti-biofilm effects against *Pseudomonas aeruginosa*"

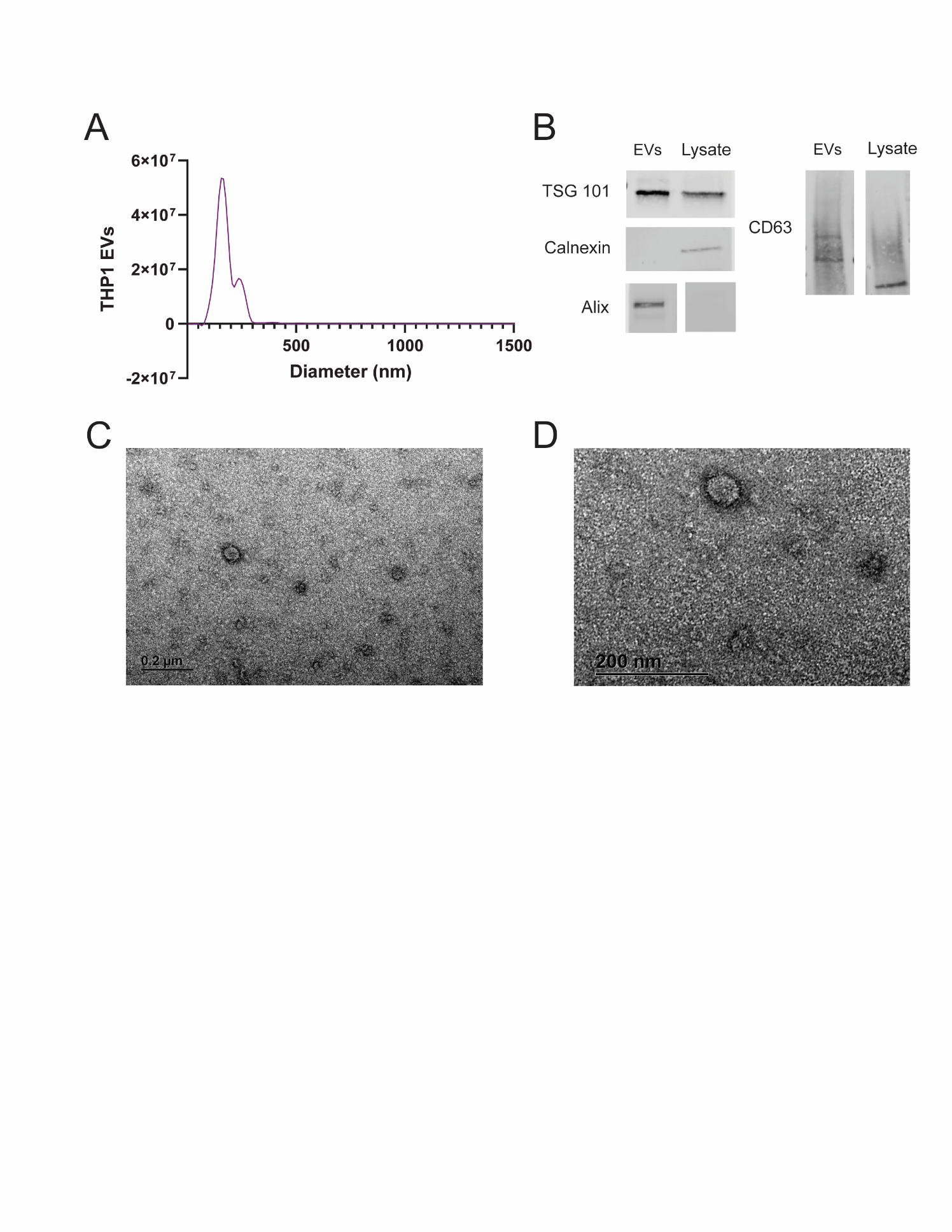

**Figure S1. THP-1 EV characterization.** THP-1 EV size distribution was determined by nanoparticle tracking analysis (NTA) (A). Western blotting for THP-1 EV samples and THP-1 cell lysate for EV markers: TSG101, CD63, and Alix as well as intracellular marker calnexin (B). Western blot data and EV size distribution are representative of at least 3 biological replicates, meaning at least 3 EV batches. TEM images of THP-1 EVs post isolation (C, D).

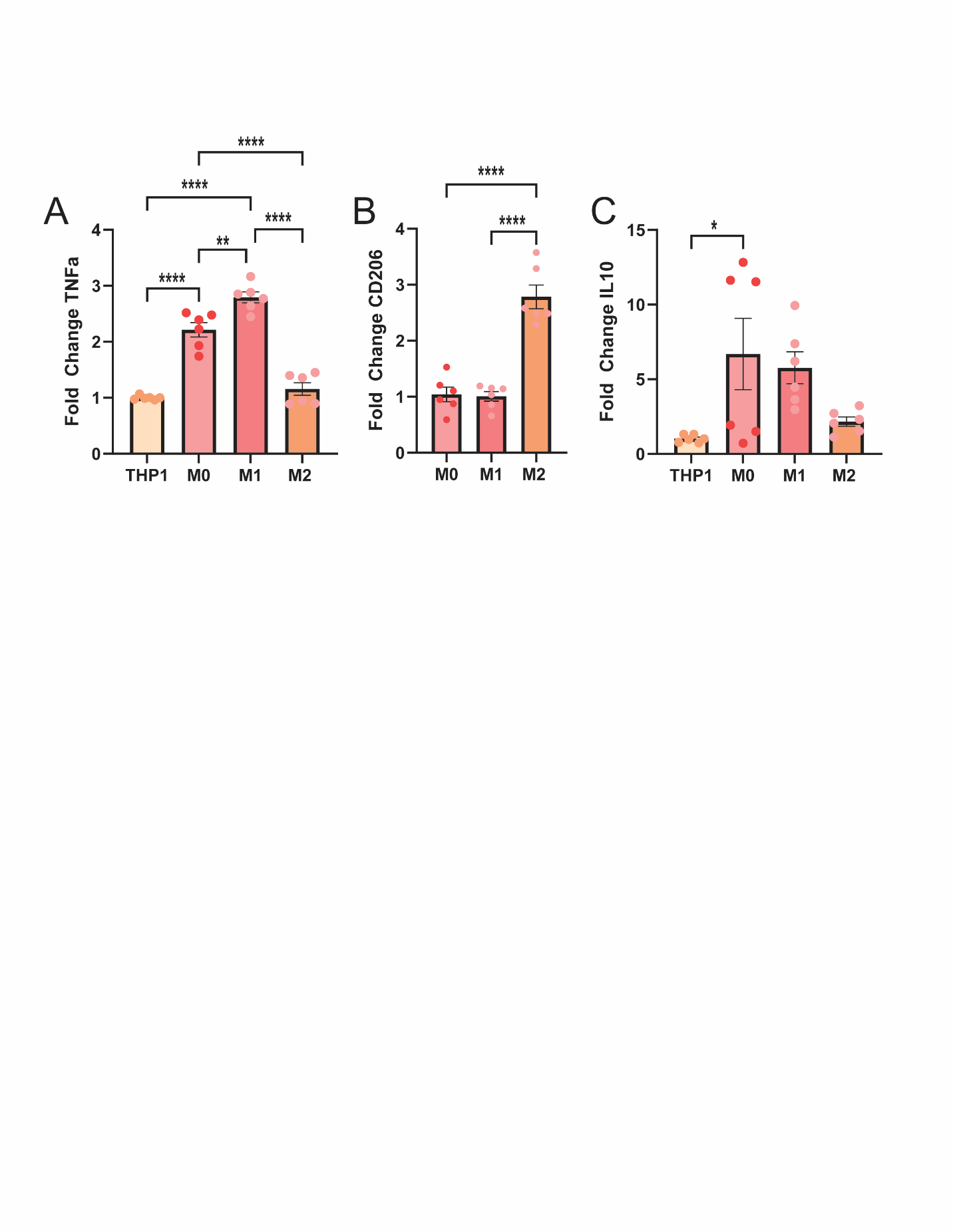

**Supplemental Figure 2. THP-1 cells are properly differentiated into macrophages and are polarized.** qPCR results for fold change in expression of TNFa (A), CD206 (B), and IL10 (C). Results are normalized to THP-1 level expression for TNFa and IL10. CD206 expression was normalized to M0 expression since THP-1 expression was indeterminate. All gene expressions were normalized to GAPDH expression. Results are two combined biological replicates.
